## Supplementary Information for "Evolutionary origin of terpenoid biosynthesis in termites"

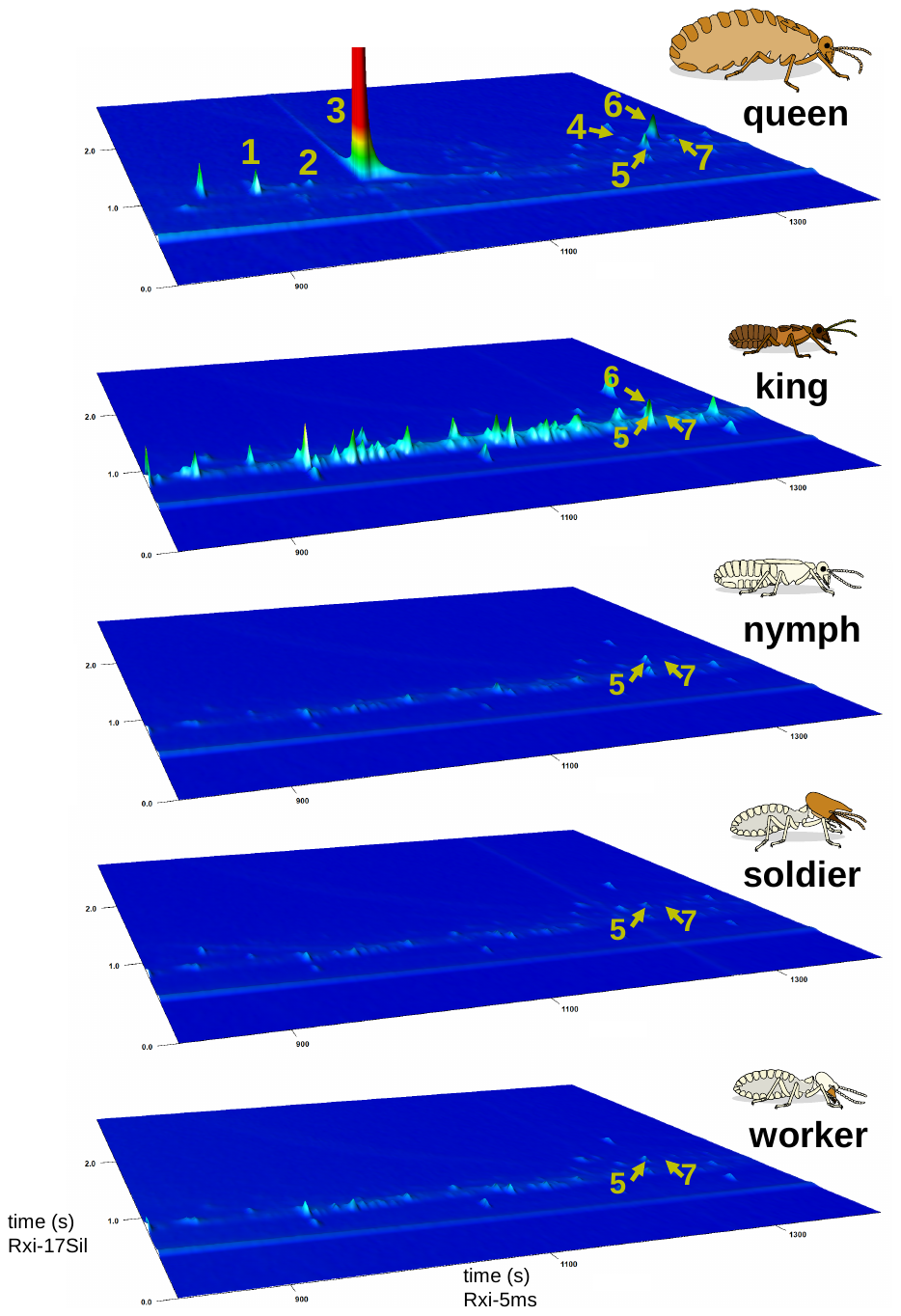

**Supplementary Fig. S1.** **Representative GC×GC chromatograms of body washes from different *E. neotenicus* castes.** The chromatograms show a retention window where terpenoids were detected. 1, β-farnesene; 2, α-farnesene; 3, (6*E*)-nerolidol; 4, unidentified diterpene; 5, β-springene; 6, (*E*,*E*,*E*)-neocemberene, 7, unidentified diterpene. The chromatogram visualizes the responses to diagnostic ions for mono-, sesqui- and diterpenes (*m/z* = 68 + 69 + 93 + 204 + 272). Mass spectra of the identified terpenes are shown below in Supplementary Fig. S2.

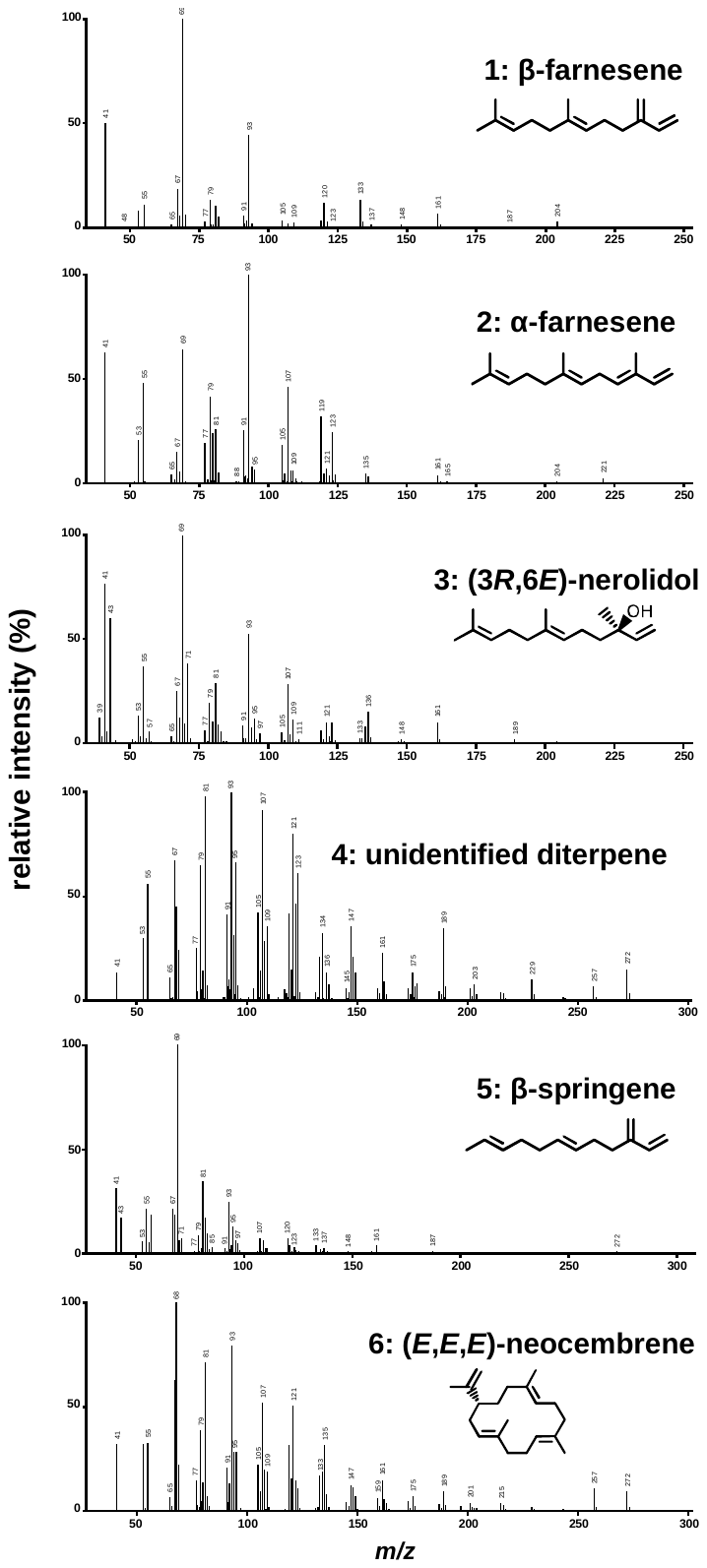

**Supplementary Fig. S2.** **Mass spectra of the terpenes identified in the body washes of different *E. neotenicus* castes.** Compound numbering refers to that in Fig. 1 and Supplementary Fig. S1. For compound 7 (unidentified diterpene), we could not obtain a reliable mass spectrum.

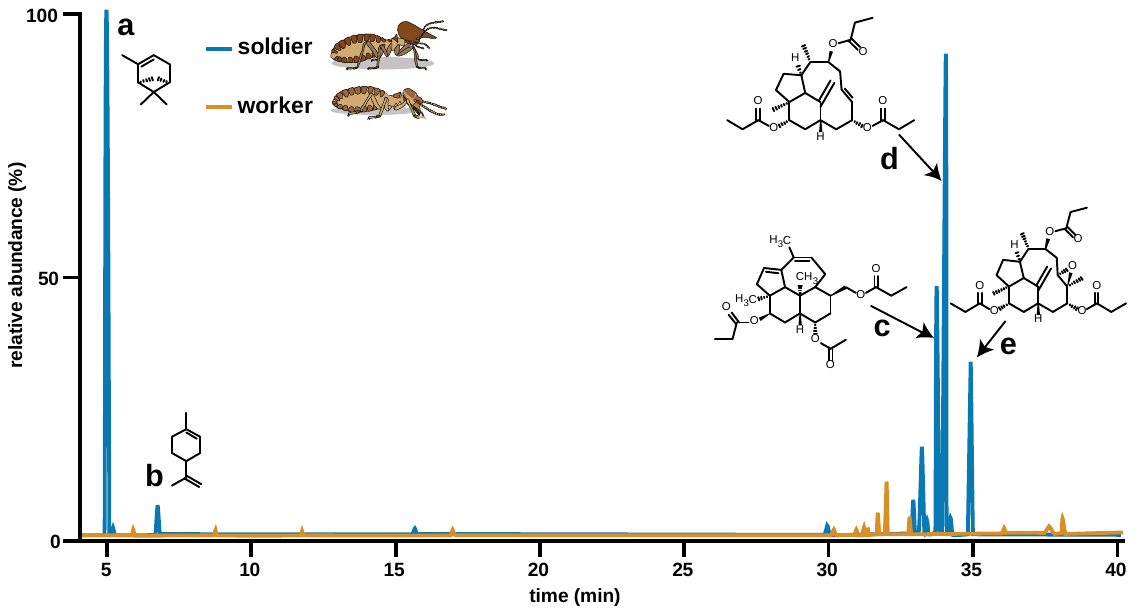

**Supplementary Fig. S3.** **Representative GC chromatogram of body washes from *N. takasagoensis* soldiers and workers.** The chromatogram visualizes the responses to diagnostic ions for mono-, sesqui- and diterpenes (*m/z* = 68 + 69 + 93 + 204 + 272). Mass spectra and NMR spectra of the identified terpenes are shown below in Supplementary Figs S4–S8.

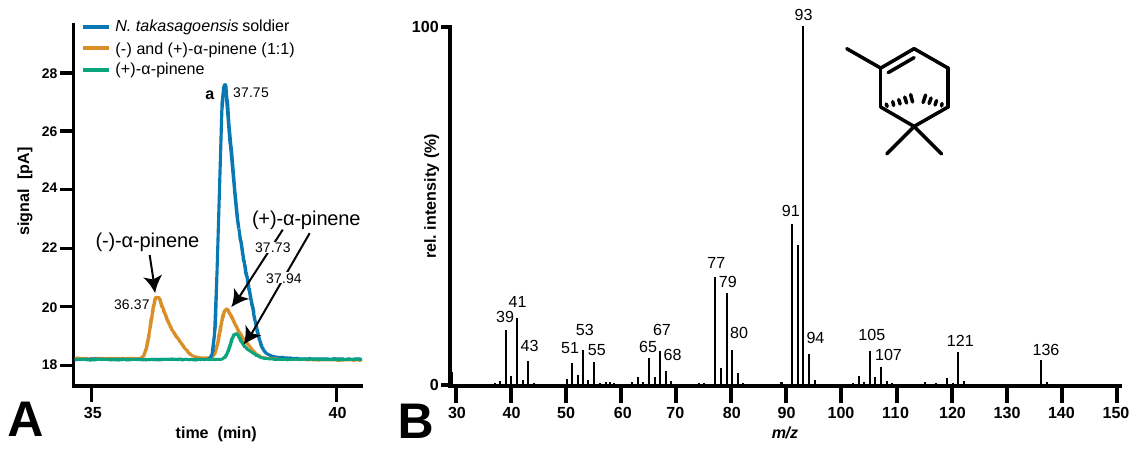

**Supplementary Fig. S4.** **Identification of compound a from *N. takasagoensis* soldiers as (+)-α-pinene**. **A.** Chiral chromatogram comparing the retention of compound **a** with standards of (-) and (+)-α-pinene. GC-FID was equipped with cyclodextrin chiral column (HP CHIRAL β, 30m, 0.25mm, 0.25µm, 19091G-B233E). One µL of sample was injected in splitless mode following this temperature program: 40 °C held for 20 minutes, ramped 1 °C/min to 100 °C, ramp at 30 °C/min to 240 °C, held for 20 minutes. **B.** Mass spectrum of compound **a**.

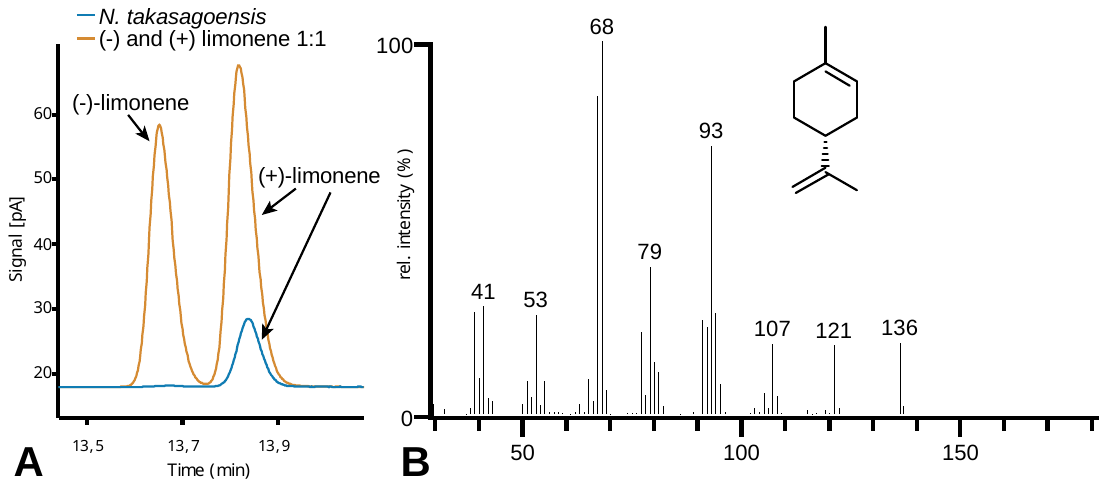

**Supplementary Fig. S5.** **Mass spectrum and chiral analysis of compound b from *N. takasagoensis* soldiers identified as (+)-limonene**.

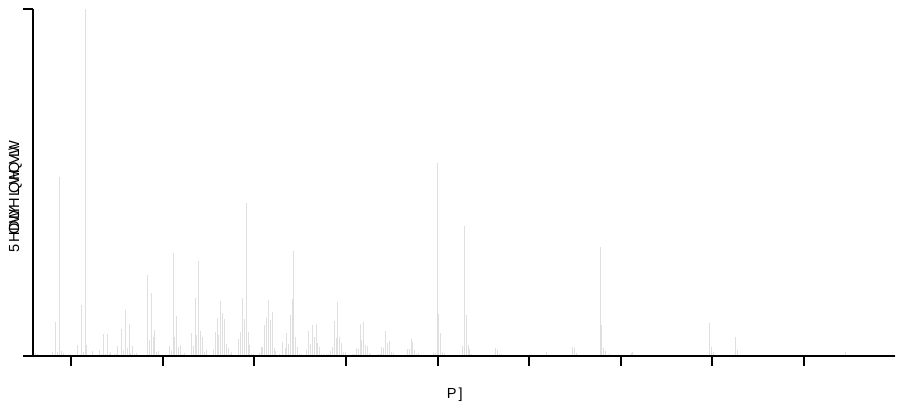

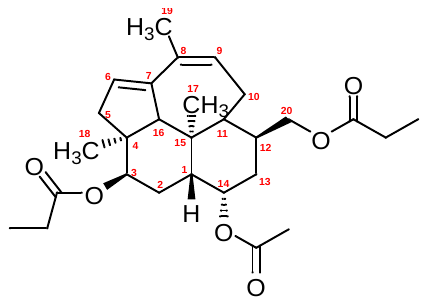
**A**

**c: 14a-acetoxy-3b,12b-dipropionoxy-kampa-6(7),8(9)-diene**

^1^H NMR (600.1 MHz, CD_2_Cl_2_) δ 5.67 – 5.64 (m, 2H, H-6,9), 5.23 (dd, *J*_3,2a_ = 9.7, *J*_3,2b_ = 7.5 Hz, 1H, H-3), 4.94 (td, *J*_14,1_ = *J*_14,13a_ = 3.3, *J*_14,13b_ = 2.5 Hz, 1H, H-14), 4.11 (dd, *J*_20a,20b_ = 11.3, *J*_20a,12_ = 5.2 Hz, 1H, H-20a), 3.91 (dd, *J*_20b,20a_ = 11.3, *J*_20b,12_ = 3.2 Hz, 1H, H-20b), 2.71 (dq, *J*_5a,5b_ = 17.8, *J*_5a,6_ = *J*_5a,18_ = 2.0, Hz, 1H, H-5a), 2.37 – 2.24 (m, 4H, CH_3_C**H_2_**CO), 2.21 (d, *J*_6,5a_ = 2.0 Hz, 1H, H-6), 2.18 (bm, 1H, H-10a), 2.11 (ddd, *J*_2a,2b_ = 14.2, *J*_2a,1_ = 11.7, *J*_2a,3_ = 9.7 Hz, 1H, H-2a), 2.05 (s, 3H, CH_3_CO), 2.04 (m, 1H, H-10b), 2.01 (m, 1H, H-10b), 1.93 (dd, *J*_5b,5a_ = 17.8, *J*_5b,6_ = 3.6 Hz, 1H, H-5b), 1.89 – 1.81 (m, 3H, H-1,12,13a), 1.81 (t, *J*_19,6_ = *J*_19,9_ = 1.6 Hz, 3H, H-19), 1.53 (m, 1H, H-11), 1.46 – 1.38 (m, 2H, H-2b,13b), 1.14 (s, 3H, H-17), 1.13 – 1.06 (m, 6H, C**H_3_**CH_2_CO), 1.00 (s, 1H, H-18).

^13^C NMR (150.9 MHz, CD_2_Cl_2_) δ 174.69 (CH_3_CH_2_**C**OO-20), 174.54 (CH_3_CH_2_**C**OO-3), 170.98 (CH_3_**C**OO-14), 144.70 (C-7), 134.21 (C-8), 130.78 (C-9), 126.37 (C-6), 74.94 (C-3), 72.02 (C-14), 68.41 (C-16), 67.09 (C-20), 51.37 (C-11), 46.08 (C-4), 41.20 (C-5), 40.57 (C-15), 39.73 (C-1), 36.39 (C-13), 32.15 (C-12), 29.73 (C-18), 28.36, 28.13 (CH_3_**C**H_2_COO), 27.54 (C-2), 25.24 (C-10), 22.79 (C-19), 21.81 (**C**H_3_CO), 20.03 (C-17), 9.64, 9.53 (**C**H_3_CH_2_COO).

**B**

**Supplementary Fig. S6, part 1. Identification of compound c from *N. takasagoensis* soldiers as 14a-acetoxy-3b,12b-dipropionoxy-kampa-6(7),8(9)-diene. A.** Mass spectrum. **B.** Assigned ^1^H and ^13^C NMR spectra.

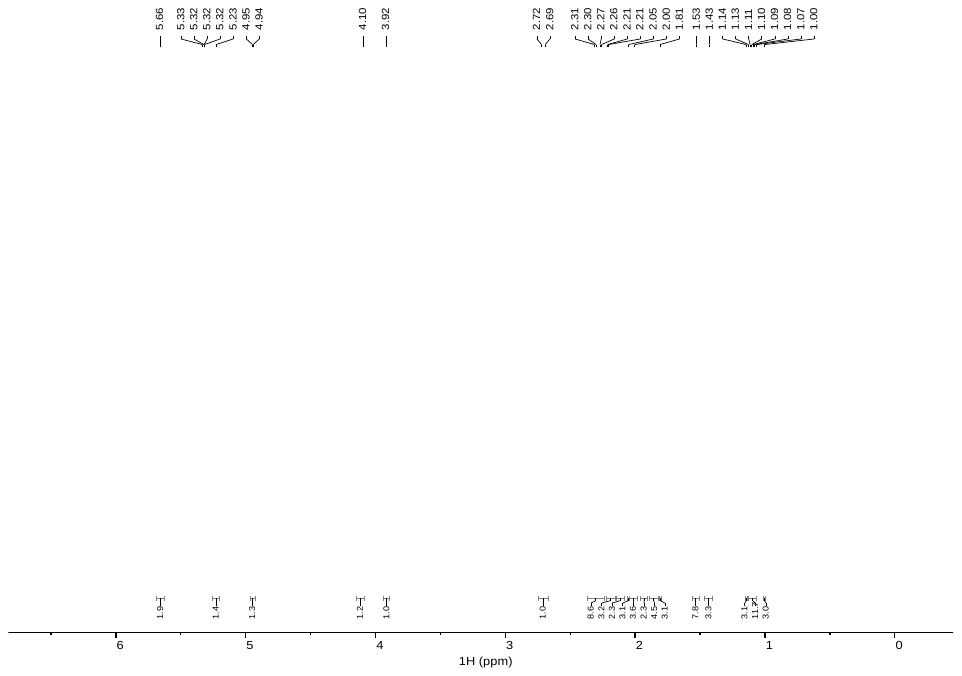

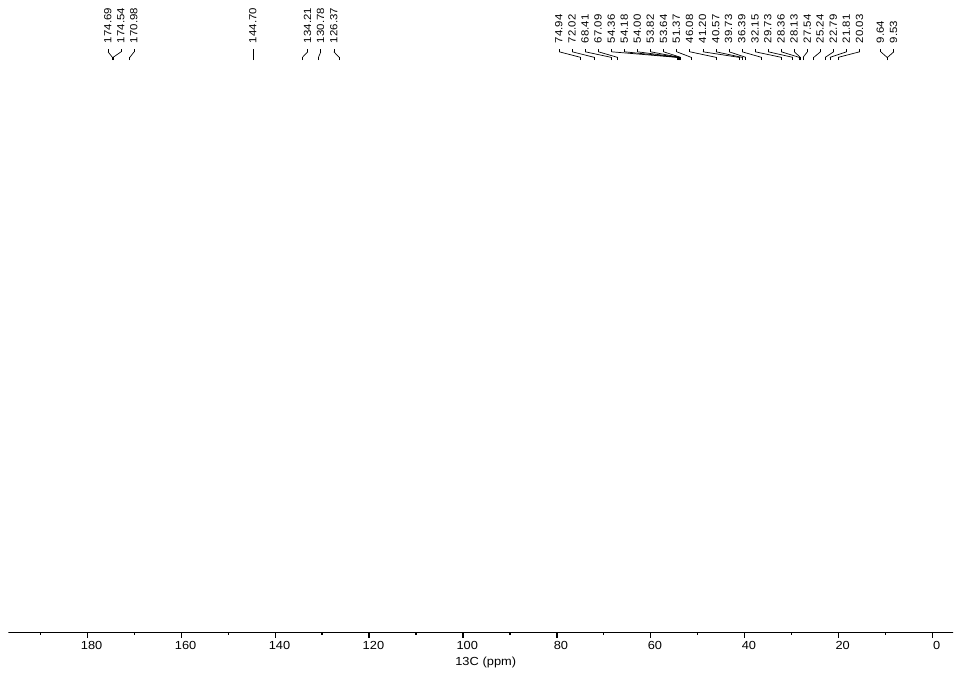
**C**

**D**

**Supplementary Fig. S6, part 1I. Identification of compound c from *N. takasagoensis* soldiers as 14a-acetoxy-3b,12b-dipropionoxy-kampa-6(7),8(9)-diene. C.** ^1^H NMR spectra. **D.** APT NMR spectrum.

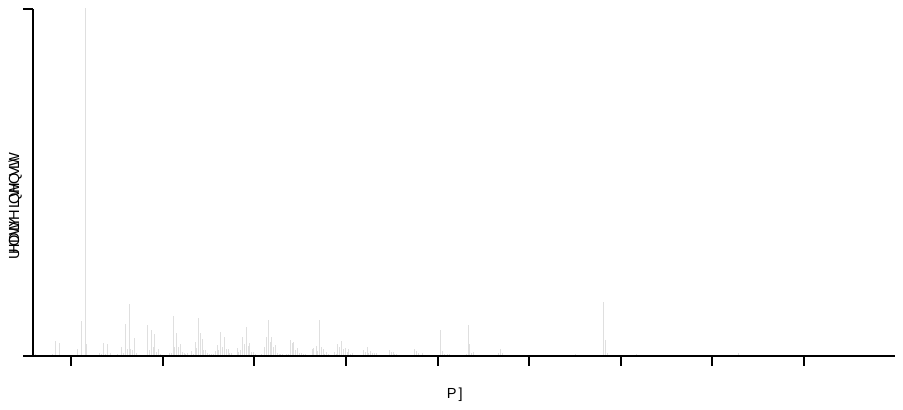

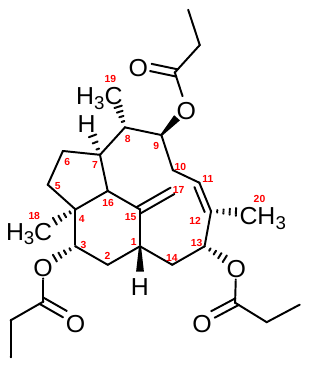
**A**

**d: 3α,9β,13α-tripropionoxy-trinervita-11(12),15(17)-diene**

^1^H NMR (600.1 MHz, CD_2_Cl_2_) δ 5.39 (m, 1H, H-11), 5.13 (dd, *J*_9,10a_ = 11.6, *J*_9,10b_ = 4.4 Hz, 1H, H-9), 5.07 (dd, *J*_13,14b_ = 11.0, *J*_13,14a_ = 6.3 Hz, 1H, H-13), 5.05 (s, 1H, H-17a), 5.03 (s, 1H, H-17b), 4.93 (dd, *J*_3,2b_ = 11.8, *J*_3,2a_ = 4.4 Hz, 1H, H-3), 2.60 (d, *J*_16,7_ = 11.4 Hz, 1H, H-16), 2.32 – 2.25 (m, 7H, H-10a, CH_3_C**H_2_**CO), 2.18 (m, 1H, H-14a), 2.08 – 2.03 (m, 2H, H-7,8), 2.01 (m, 1H, H-10b), 1.95 (m, 1H, H-1), 1.78 – 1.71 (m, 2H, H-2a,6a), 1.65 (d, *J*_20,11_ = 1.4 Hz, 3H, H-20), 1.64 – 1.53 (m, 3H, H-5a,6b,14b), 1.41 (m, 1H, H-2b), 1.15 – 1.06 (m, 10H, H-5b, C**H_3_**CH_2_CO), 0.92 (s, 1H, H-18), 0.89 (d, *J*_19,8_ = 6.2 Hz, 1H, H-19).

^13^C NMR (150.9 MHz, CD_2_Cl_2_) δ 174.58 (CH_3_CH_2_**C**OO-9), 174.54 (CH_3_CH_2_**C**OO-3), 173.87 (CH_3_CH_2_**C**OO-13), 147.90 (C-15), 136.65 (C-12), 124.46 (C-11), 114.15 (C-17), 79.97 (C-13), 74.58 (C-9), 73.69 (C-3), 60.12 (C-16), 48.76 (C-4), 44.85 (C-7), 38.96 (C-1), 38.09 (C-2), 37.38 (C-8), 36.71 (C-5), 34.10 (C-14), 30.62 (C-6), 28.72 (C-10), 28.41, 28.29 (CH_3_**C**H_2_COO), 20.10 (C-18), 12.16 (C-19), 11.89 (C-20), 9.66, 9.57, 9.53 (**C**H_3_CH_2_COO).

**B**

**Supplementary Fig. S7, part 1. Identification of compound d from *N. takasagoensis* soldiers as 3α,9β,13α-tripropionoxy-trinervita-11(12),15(17)-diene**. **A.** Mass spectrum. **B.** Assigned ^1^H and ^13^C NMR spectra.

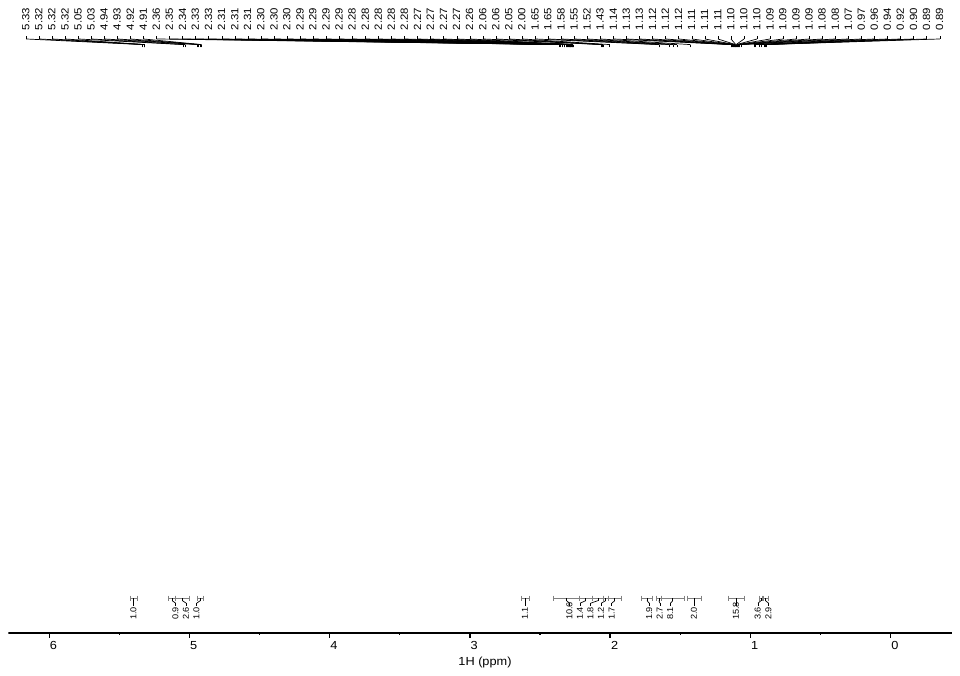

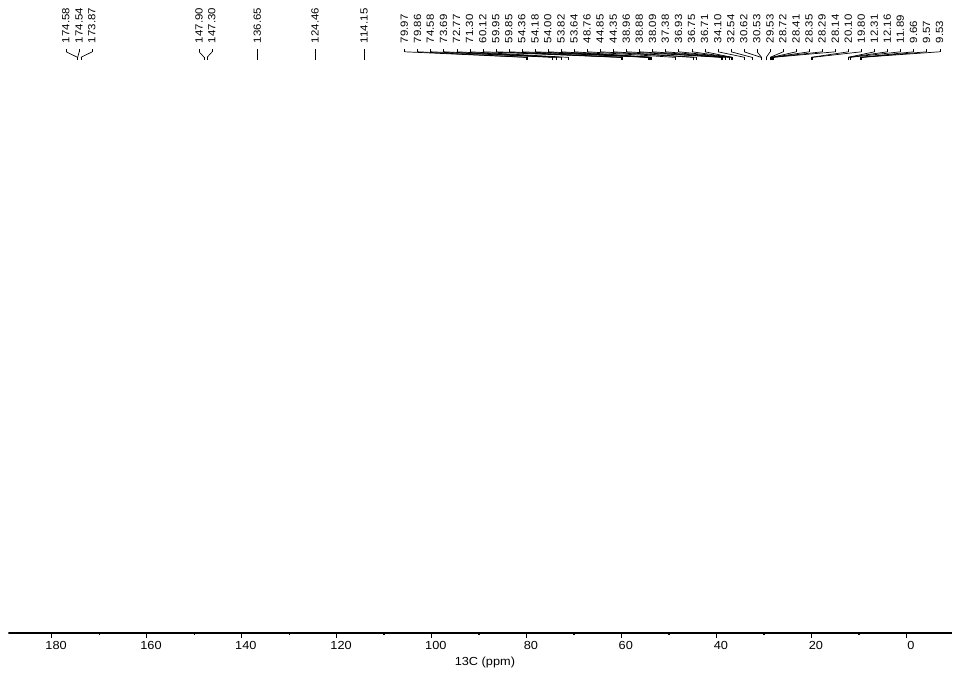

**C**

**D**

**Supplementary Fig. S7, part 1I. Identification of compound d from *N. takasagoensis* soldiers as 3α,9β,13α-tripropionoxy-trinervita-11(12),15(17)-diene**. **C.** ^1^H NMR spectra. **D.** APT NMR spectrum.

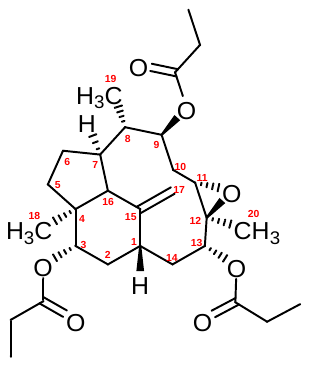

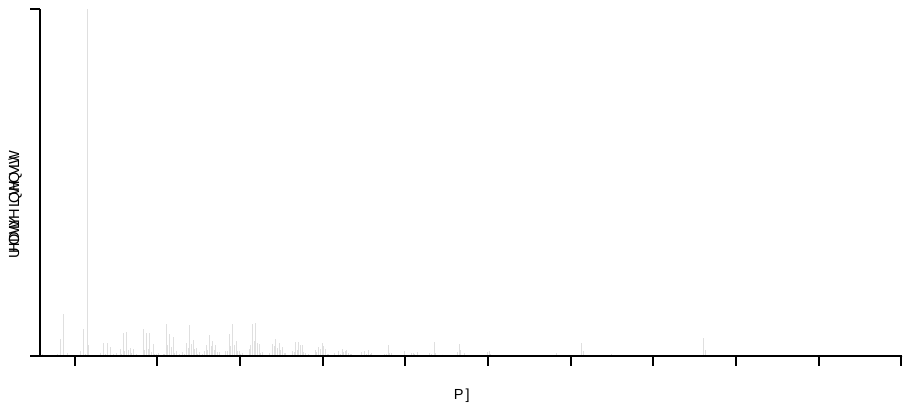
**A**

**e: 3α, 9β,13α-tripropionoxy-11α(12β)-epoxy-trinervita-15(17)-ene**

^1^H NMR (600.1 MHz, CD_2_Cl_2_) δ 5.43 (dd, *J*_9,10a_ = 11.9, *J*_9,10b_ = 3.8 Hz, 1H, H-9), 5.25 (s, 1H, H-17a), 5.16 (s, 1H, H-17b), 4.96 (dd, *J*_3,2b_ = 11.8, *J*_3,2a_ = 4.5 Hz, 1H, H-3), 4.31 (dd, *J*_13,14a_ = 11.8, *J*_13,14b_ = 5.6 Hz, 1H, H-13), 2.89 (dd, *J*_11,10b_ = 10.6, *J*_11,10a_ = 4.3 Hz, 1H, H-11), 2.61 (d, *J*_16,7_ = 11.7 Hz, 1H, H-16), 2.42 – 2.26 (m, 6H, CH_3_C**H_2_**CO), 2.26 – 2.19 (m, 2H, H-10a,14a), 2.14 (m, 1H, H-7), 2.08 (m, 1H, H-1), 1.99 (qd, *J*_8,19_ = 6.6, *J*_8,7_ = 5.4 Hz, 1H, H-8), 1.89 – 1.76 (m, 2H, H-2a,6a), 1.63 (m, 1H, H-5a), 1.59 (m, 1H, H-6b), 1.63 (m, 1H, H-5a), 1.53 (m, 1H, H-14b), 1.43 (s, 3H, H-20), 1.43 (m, 1H, H-2b), 1.30 (ddd, *J*_10b,10a_ = 12.7, *J*_10b,11_ = 10.6, *J*_10b,9_ = 3.8 Hz, 1H, H-10b), 1.17 – 1.06 (m, 10H, H-5b, C**H_3_**CH_2_CO), 0.97 (d, *J*_19,8_ = 6.6 Hz, 1H, H-19), 0.94 (s, 1H, H-18).

^13^C NMR (150.9 MHz, CD_2_Cl_2_) δ 174.49 (CH_3_CH_2_**C**OO-3), 174.20 (CH_3_CH_2_**C**OO-9), 173.82 (CH_3_CH_2_**C**OO-13), 147.30 (C-15), 114.13 (C-17), 79.86 (C-13), 72.77 (C-3), 71.31 (C-9), 59.95 (C-16), 59.85 (C-11), 59.82 (C-12), 48.77 (C-4), 44.35 (C-7), 38.96 (C-8), 38.88 (C-2), 36.93 (C-1), 36.75 (C-5), 32.54 (C-14), 30.53 (C-6), 29.53 (C-10), 28.35, 28.27, 28.14 (CH_3_**C**H_2_COO), 19.80 (C-18), 12.54 (C-20), 12.31 (C-19), 9.66, 9.54, 9.45 (**C**H_3_CH_2_COO).

**B**

**Supplementary Fig. S8, part 1. Identification of compound e from *N. takasagoensis* soldiers as 3α, 9β,13α-tripropionoxy-11α(12β)-epoxy-trinervita-15(17)-ene**. **A.** Mass spectrum. **B.** Assigned ^1^H and ^13^C NMR spectra.

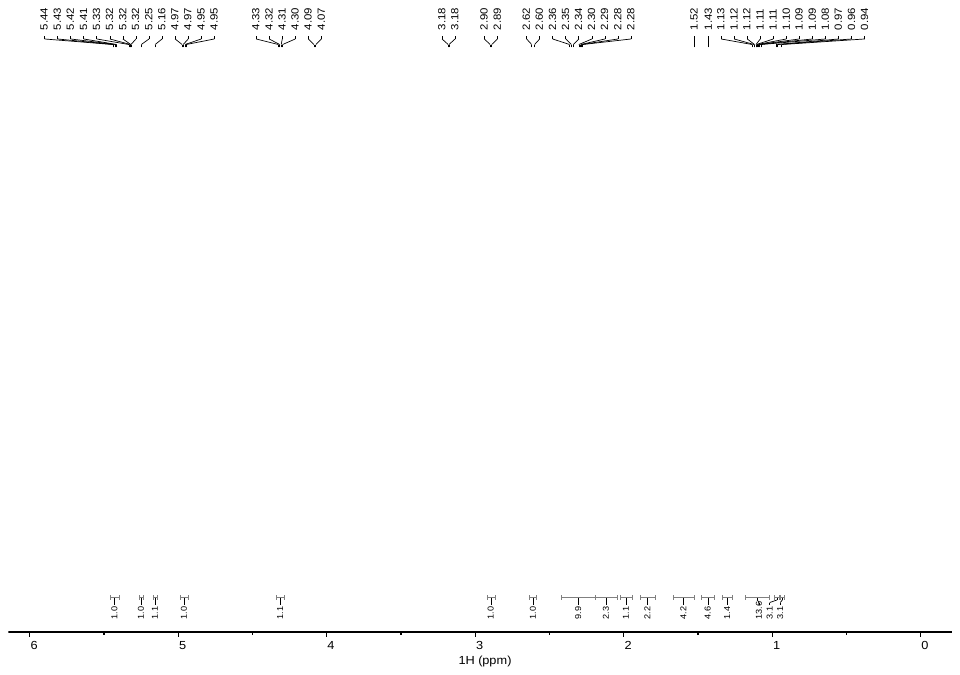

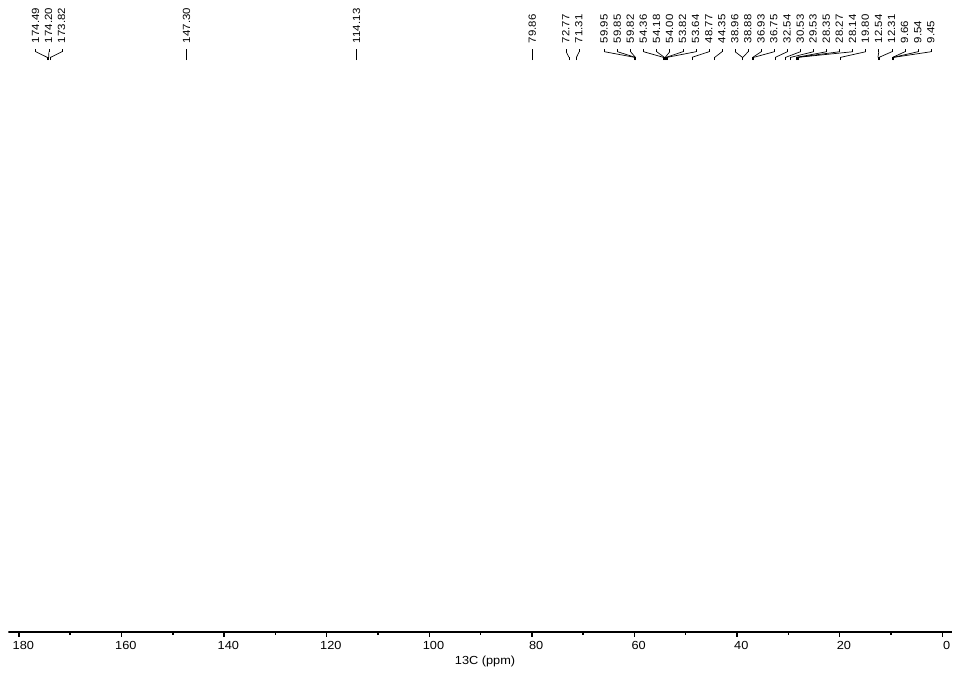
**C**

**D**

**Supplementary Fig. S8, part 1I. Identification of compound e from *N. takasagoensis* soldiers as 3α, 9β,13α-tripropionoxy-11α(12β)-epoxy-trinervita-15(17)-ene**. **C.** ^1^H NMR spectra. **D.** APT NMR spectrum.

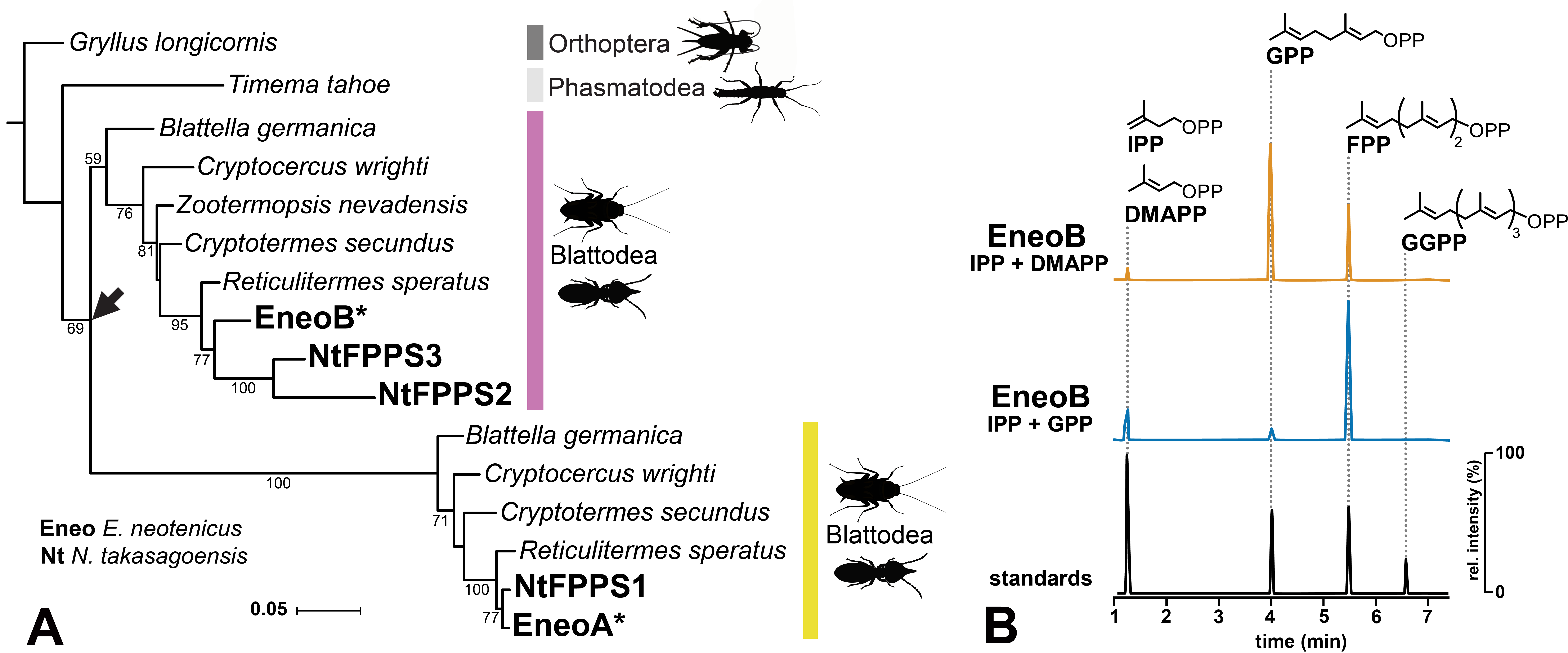

**Supplementary Fig. S9.** **Phylogeny and function of FPPS homologs in termites**. **A.** Phylogenetic tree inferred from amino acid sequences of FPPS homologs identified in Blattodea (five termite and two cockroach species) and two other polyneopteran orders, Orthoptera and Phasmatodea. The maximum likelihood tree was reconstructed with IQ-TREE 2 using JTT+G4 model; bootstrap values (500 replicates) greater than 50 are shown as node support. Arrow marks the FPPS duplication predating the diversification of Blattodea. Asterisks highlight the two paralogous sequences EneoA and EneoB from *Embiratermes neotenicus* studied with respect to their function. Accession numbers are provided in Supplementary Table S2. **B.** HPLC chromatogram demonstrating the IDS activity of EneoB. The purified enzyme was incubated with isopentenyl pyrophosphate (IPP) and dimethylallyl pyrophosphate (DMAPP), IPP and geranyl pyrophosphate (GPP), or IPP and farnesyl pyrophosphate (FPP). Only the substrate combinations giving rise to any prenyl pyrophosphate products are shown. The chromatograms visualize the selected m/z 245.00, 313.06, 381.12, and 449.19. No TPS activity was recorded in EneoB and no TPS or IDS activity was observed in EneoA.

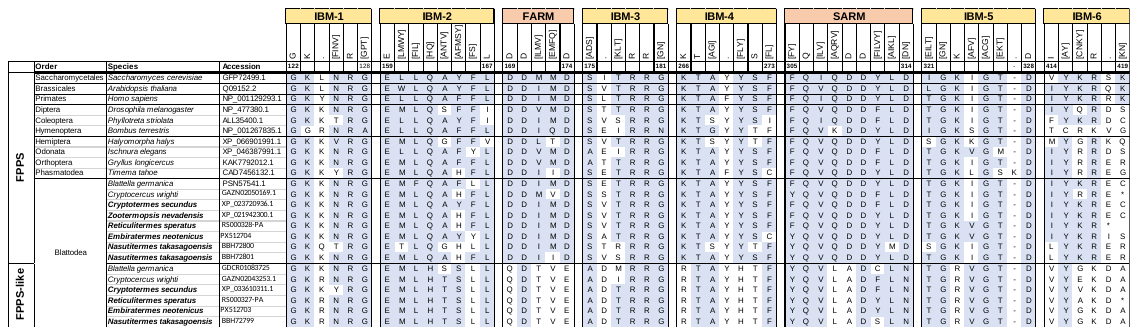

**Fig. S9. Alignment of IDS catalytic motifs in FPPS and FPPS-like amino acid sequences. Motif b**oundaries and sequence regular expressions are displayed as defined by Rebholz et al. (2023) for insect IDS sequences. Residues in consensus with the regular expression are shaded blue. Termite species are displayed in bold text. IBM, IPP-binding motif. FARM, first aspartate-rich motif. SARM, second aspartate-rich motif.

**Supplementary Fig. S10. Alignment of IDS catalytic motifs in FPPS and FPPS-like amino acid sequences. Motif b**oundaries and sequence regular expressions are displayed as defined by Rebholz et al. (2023) for insect IDS sequences. Residues in consensus with the regular expression are shaded blue. Termite species are displayed in bold text. IBM, IPP-binding motif. FARM, first aspartate-rich motif. SARM, second aspartate-rich motif.

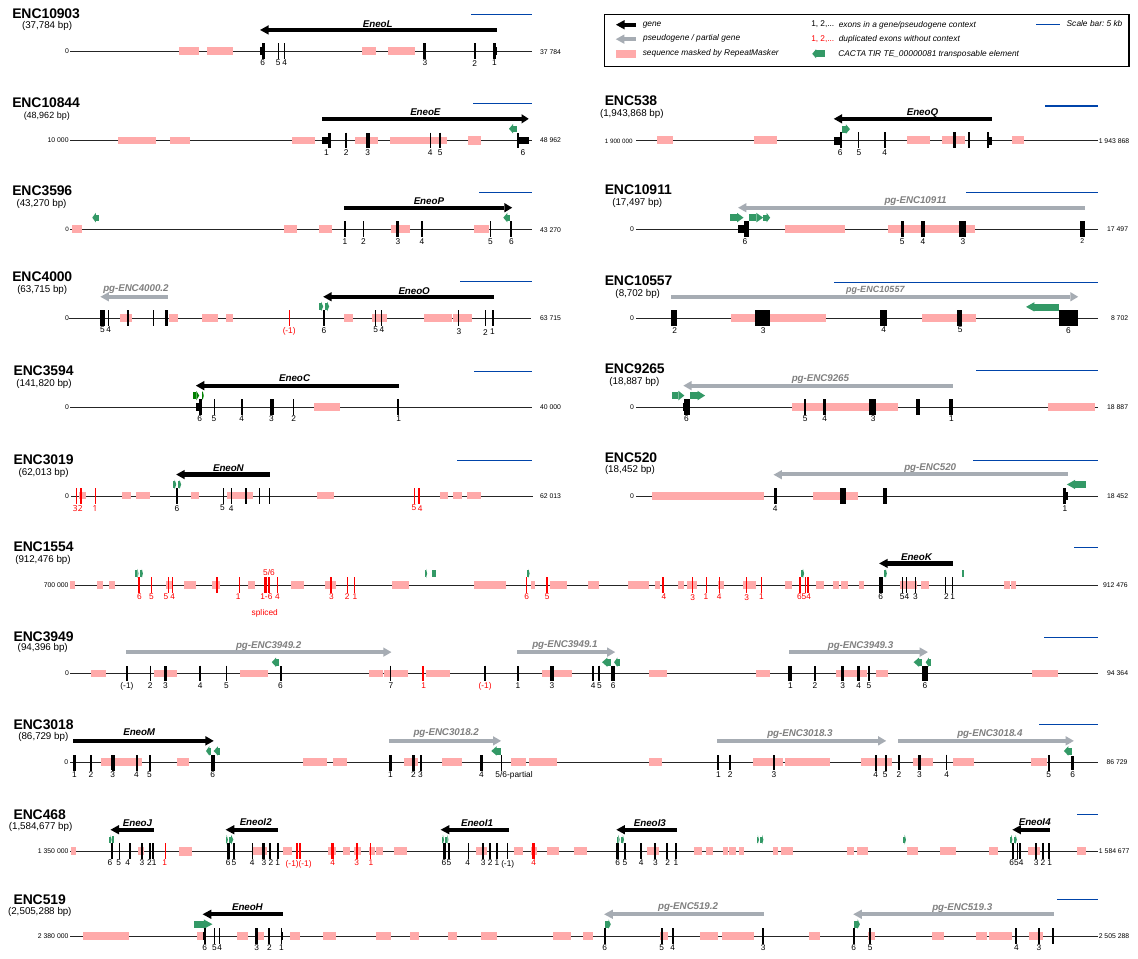

**Supplementary Fig. S11. Genomic landscape surrounding the *E. neotenicus* GGPPS-like genes and pseudogenes.** The figure shows gene organization of full or partial genomic contigs (boldfaced, starting with prefix ENC and contig size in brackets) represented by horizontal line with position coordinates at both sides. Exons are numbered and shown as black (genes), grey (pseudogenes) or red boxes (chimeric duplicates), regions corresponding to genes and pseudogenes are marked with black or grey arrows while regions masked by RepeatMasker at genome-wide scale are shown as pink boxes. Sequences homologous to CACTA TIR TE_00000081 DNA transposon predicted with the EDTA pipeline are marked with green arrows. Blue scale bar represents 5 kb.

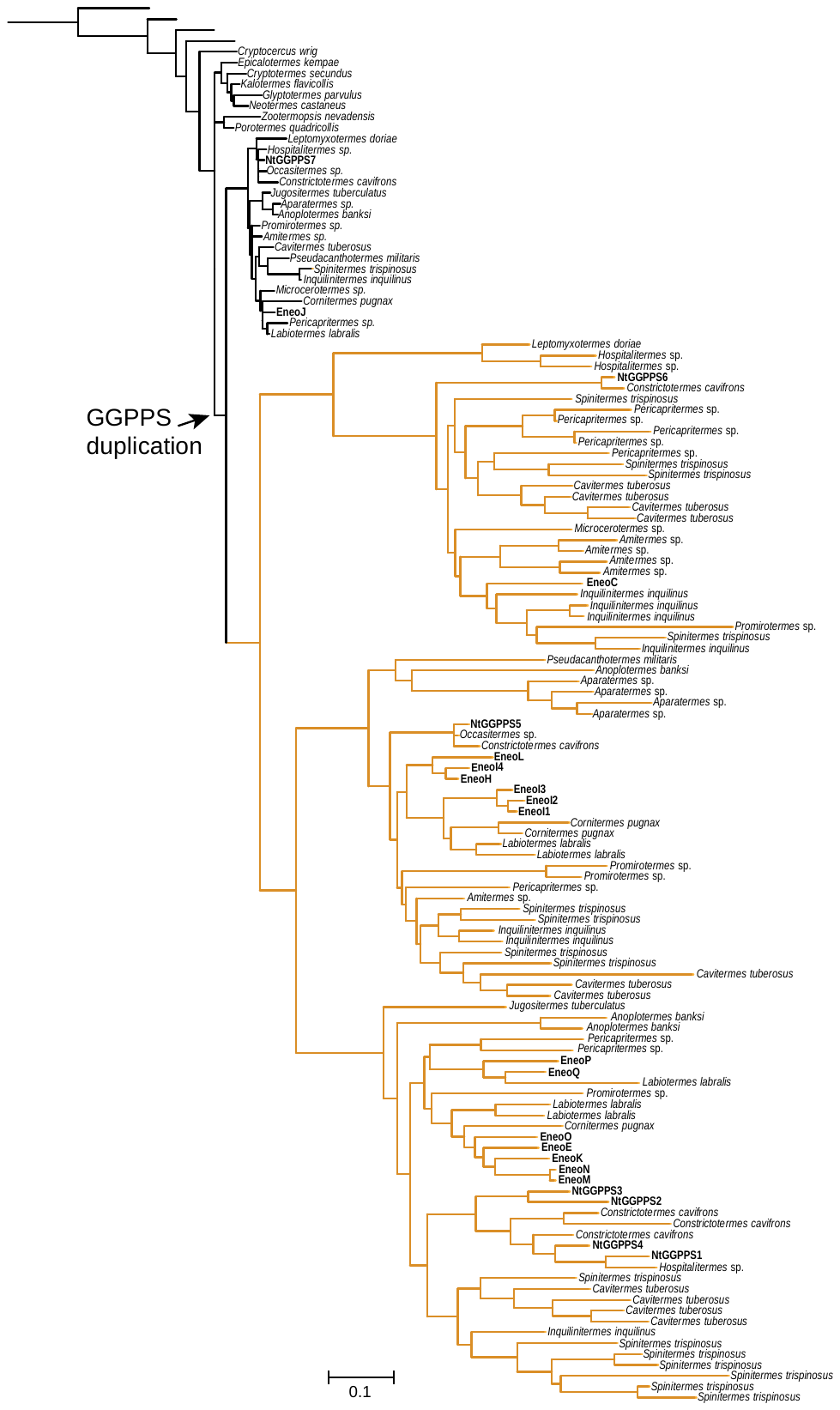
**Supplementary Fig. S12. Maximum-likelihood phylogenetic tree inferred from alignment of Blattodea GGPPS amino acid sequences.** The tree was reconstructed using IQ-TREE with the JTT+F+R4 substitution model; bootstrap values (1,000 replicates) greater than 50 are shown as node support. The tree was rooted with a dragonfly GGPPS sequence (*Ischnura elegans*). See Supplementary Table S5 for source sequences.

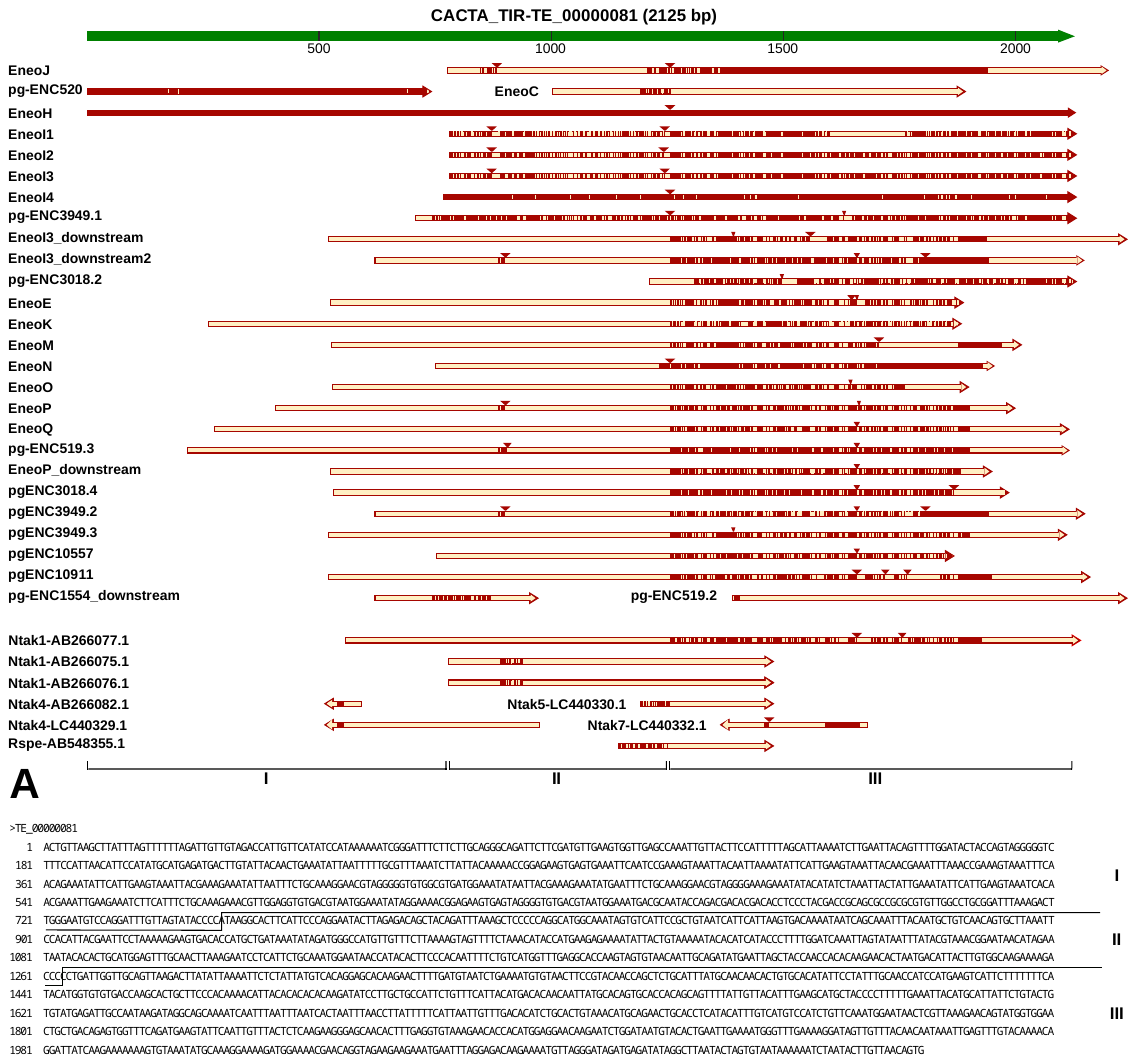
 **Supplementary Fig. S13. Sequence conservation of the CACTA TIR DNA transposable element likely involved in GGPS-like gene multiplication in Neoisoptera.** **A.** Sequences corresponding to CACTA TIR TE_00000081 predicted with RepeatModeler (green) and homologous sequences found in the proximity of GGPPS-like genes and pseudogenes in *E. neotenicus* (yellow) are shown as arrows, conserved regions are represented by dark color. Red triangles above sequence diagrams represent indels relative to the CACTA TIR TE_00000081 reference. Orthologs from related species *N. takasagoensis* (Ntak) and *R. speratus* (Rspe) found by BLAST search against the complete NCBI nucleotide collection restricted to Blattodea are included. **B.** The sequence of CACTA TIR TE_00000081 predicted with RepeatModeler, latin numbers I-III are used to specify three regions with various levels of conservation.

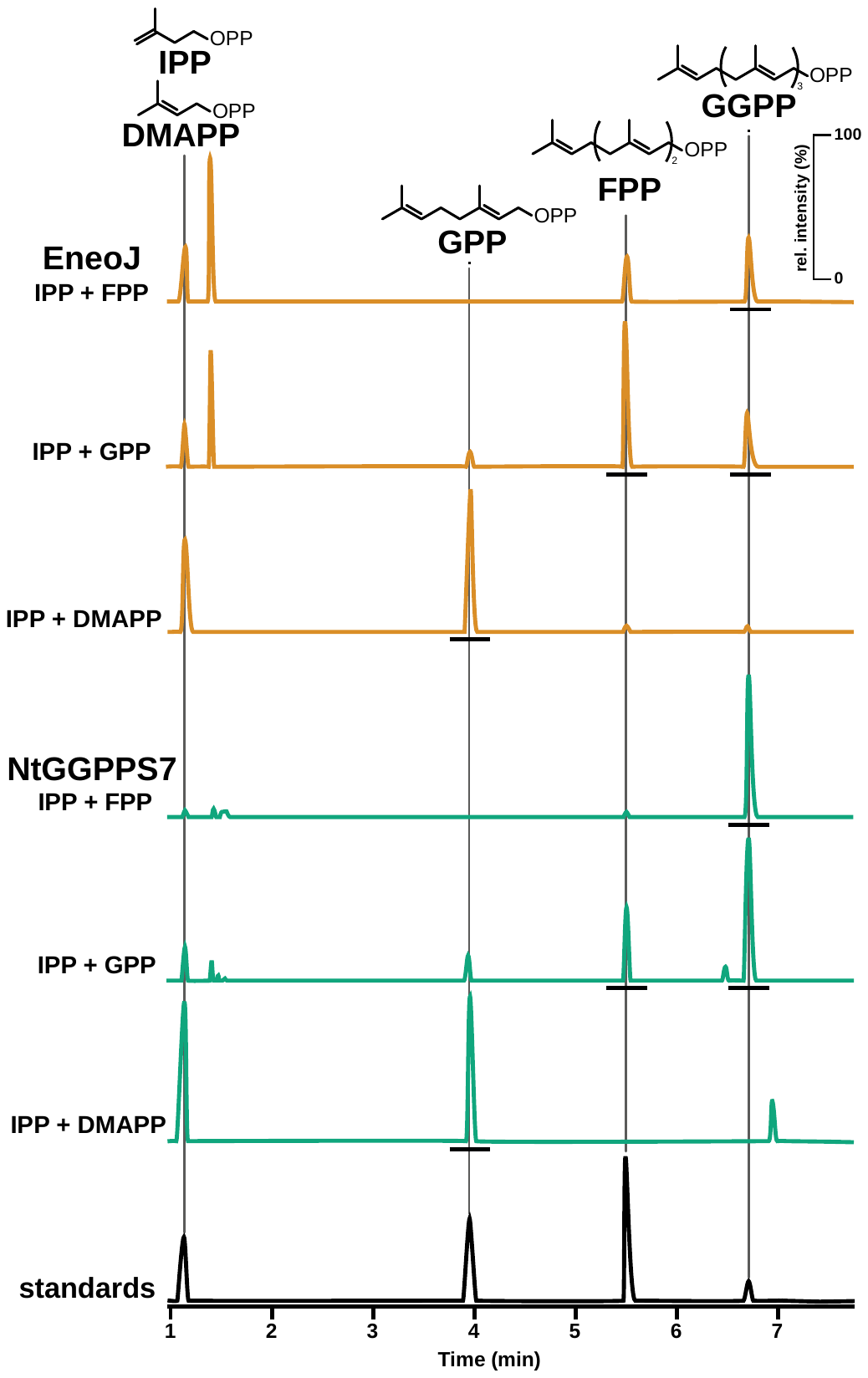

**Supplementary Fig. S14.** **HPLC chromatograms demonstrating the IDS activity in canonical GGPPSs EneoJ (yellow) and NtGGPPS7 (green).** The purified enzymes were incubated with IPP and DMAPP, IPP and GPP, or IPP and FPP. The chromatograms visualize the selected m/z 245.00, 313.06, 381.12, and 449.19. The products resulting from IDS activity are underlined with black lines.

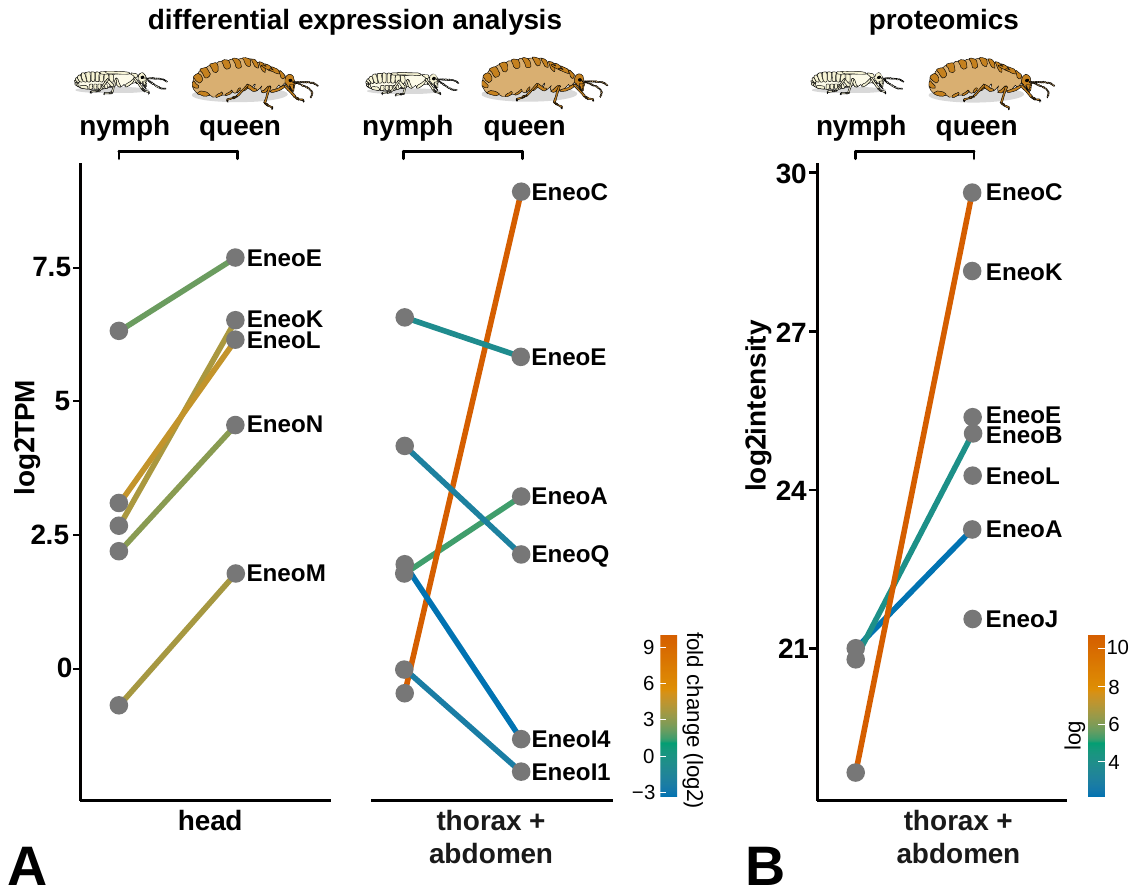

**Supplementary Fig. S15. Expression of IDS and IDS-like genes in *E. neotenicus* queens and nymphs at the level of transcript (A) and protein (B). A.** Differential expression analysis comparing transcript abundances of IDS and IDS-like genes in heads or thoraces and abdomens of neotenic queens and queen-destined 4^th^ stage female nymphs. Mean log_2_ TPM values represent three biological replicates, colors of connecting lines correspond to log_2_ fold changes between the two castes. Only genes which showed significant regulation between the two castes in DESEq analysis are presented (padj < 0.05). Expression data for all IDS/IDS-like genes are listed in Supplementary Table S7. **B.** Quantitative proteomic analysis of IDS/IDS-like proteins in thoraces and abdomens of queen-destined 4^th^ stage female nymphs and neotenic queens. Mean log_2_ intensity values represent four biological replicates. Colors correspond to log_2_ fold changes between the two castes. Only proteins which conform to the permutation-based FDR 0.05 are shown; points with no connecting lines represent proteins which were only detected in samples from queen thoraces and abdomens. Results of proteomic analysis for all IDS/IDS-like proteins from all tissues are listed in Supplementary Table S8.

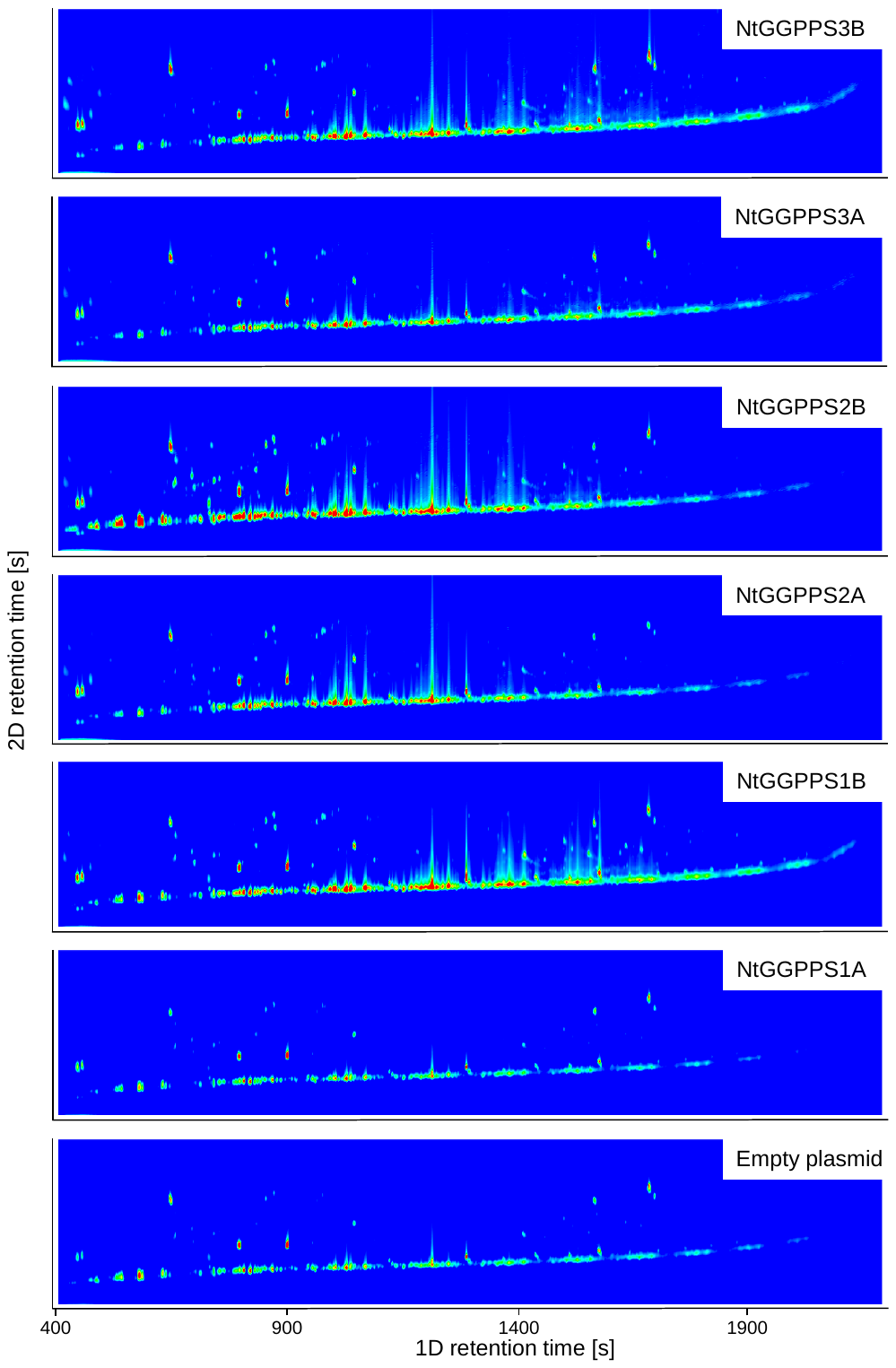

**Supplementary Fig. S16, part 1. Screening of enzymatic activity of GGPPS-like enzymes from *Nasutitermes takasagoensis.***

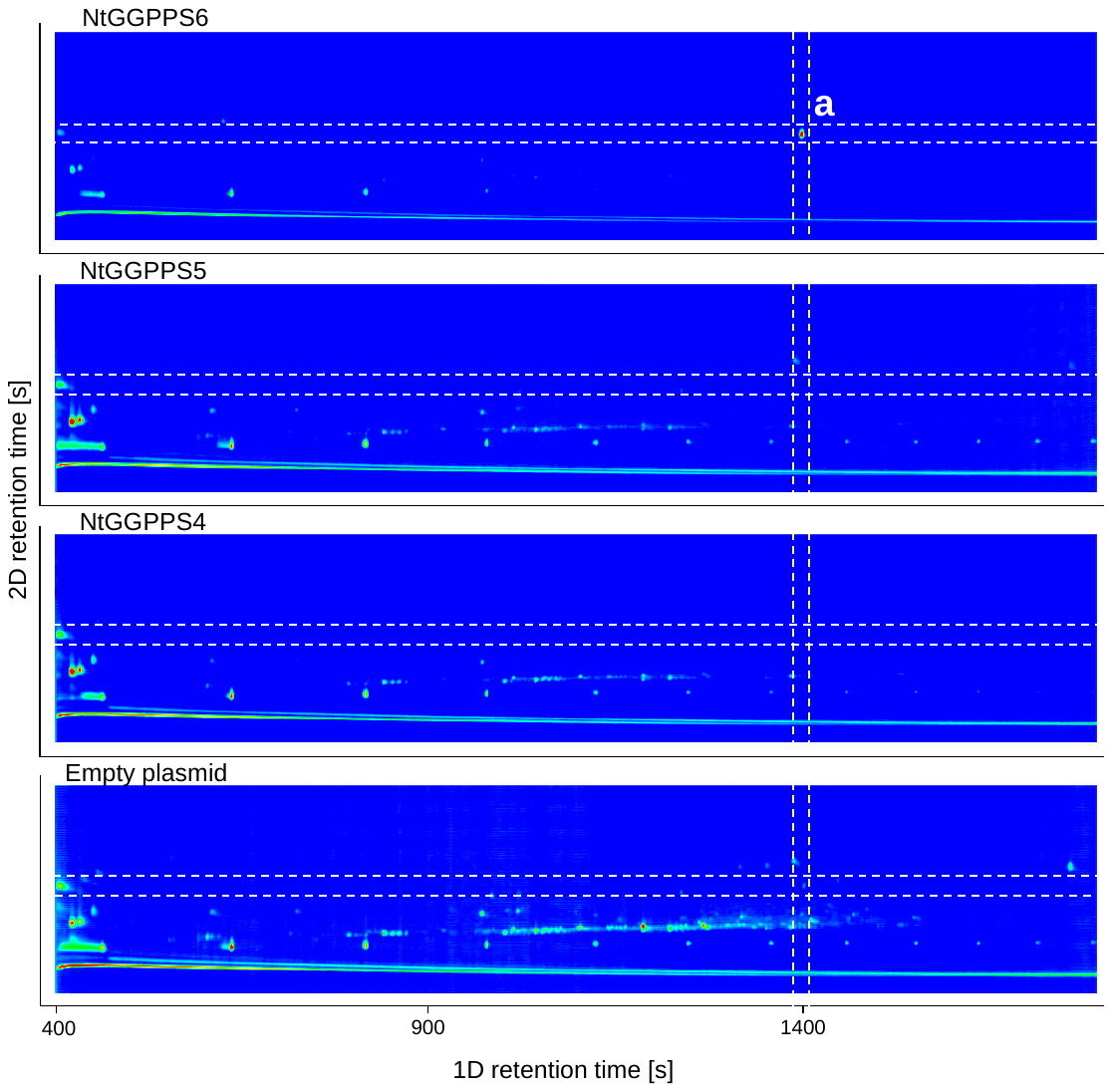
**Supplementary Fig. S16, part II. Screening of enzymatic activity of GGPPS-like enzymes from *Nasutitermes takasagoensis.*** Sequences were expressed in *S. cerevisiae* strain JWY501. 1mL of culture was extracted with 1mL of hexane and 1 μL aliquot of hexane extract was measured by comprehensive gas chromatography mass spectrometry. No specific products were detected in any of the strains apart from neocembrene (a) in the NtGGPPS6-expressing strain.

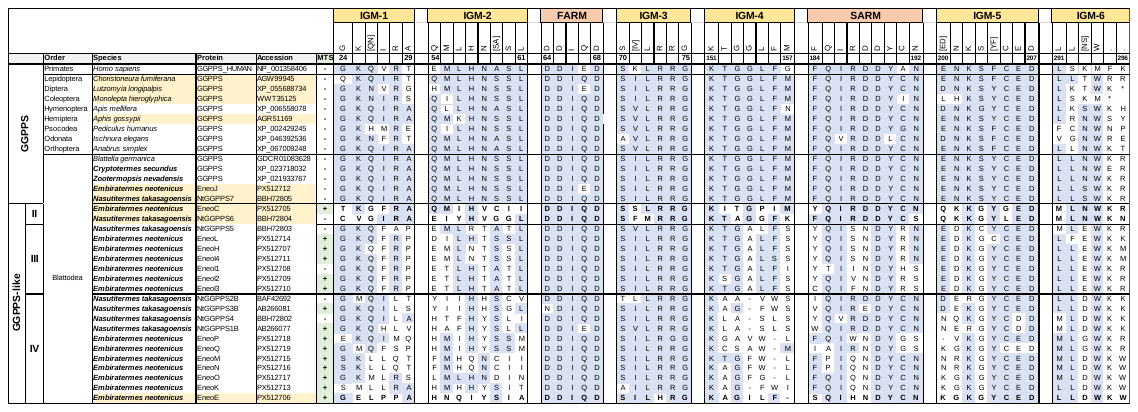

**Supplementary Fig. S17.** **Alignment of IDS catalytic motifs in GGPPS and GGPPS-like protein sequences.** Boundaries of motifs are displayed as defined by Rebholz et al. (2023). Insect GGPPS-specific regular expression for each motif was generated based on the canonical insect GGPPS sequences displayed in the alignment using MEME (v5.5.7). Residues in consensus with the regular expression are shaded blue. Termite species are printed in bold text, amino acid residues for confirmed TPS proteins are set in bold. MTS, presence or absence of a mitochondrial targeting peptide as predicted by MitoFates (v1.2). IGM, insect GGPPS motif. FARM, first aspartate-rich motif. SARM, second aspartate-rich motif.

Rebholz Z, Shewade L, Kaler K, Larose H, Schubot F, Tholl D, et al. Emergence of terpene chemical communication in insects: Evolutionary recruitment of isoprenoid metabolism. Protein Science. 2023; 32(5):e4634. <https://doi.org/10.1002/pro.4634>
